# Kronos: A computational tool to facilitate biological rhythmicity analysis

**DOI:** 10.1101/2023.04.21.537503

**Authors:** Thomaz F. S. Bastiaanssen, Sarah-Jane Leigh, Gabriel S. S. Tofani, Cassandra E. Gheorghe, Gerard Clarke, John F. Cryan

## Abstract

**Motivation:** Circadian rhythms, or 24-hour biological cycles, are key in maintaining health in almost all living organisms and synchronize important physiological and behavioural processes daily. Interest in circadian rhythm research is expanding as our urban environments have increased exposure to factors that can disrupt the normal physiological rhythm of our body, such as delayed bedtimes, shift work, jet-lag, increased screen-time, and exposure to artificial light. Discovering how oscillatory signals respond to both external and internal factors can lead to important biological breakthroughs, but assessing rhythmicity can be both limited by the complexity of statistical models and demanding in terms of coding and statistical expertise.

**Implementation:** Here, we describe the development of a novel easy-to-use R-based tool, Kronos, to assess circadian rhythms in biological data sets. Kronos provides the user with new functionalities not currently available, including the analysis of two or more groups in complex study designs, handling both independent and repeated-measures data, as well as ranging from single variables to high dimensional ‘omics data sets. Kronos is a novel tool to facilitate the analysis of rhythmicity in simple and complex experimental designs and enables researchers from diverse scientific fields to interrogate rhythmicity.

**Availability:** https://github.com/thomazbastiaanssen/kronos

## 1 Introduction

### 1.1 Circadian Rhythms

Chronobiology is defined as the study of the effect of time on biological rhythms (Caliyurt, 2017). All biological processes of living organisms, from humans to microorganisms (Loudon, 2012) and indeed the interplay between them (Teichman et al., 2020), are regulated by a 24-hour cycle referred to as circadian rhythm. Circadian rhythms are investigated in multiple fields, having metabolic (Marcheva et al., 2013), immune (Scheiermann et al., 2013), and mental and behavioural (Walker et al., 2020) implications. In mammals, circadian rhythms are coordinated by our master clock located in the hypothalamus - more specifically in the suprachiasmatic nucleus (Jennifer et al., 2012). This tightly regulated system is controlled by a complex set of feedback loop mechanisms to regulate sleep/wake cycle, body temperature, energy metabolism and even behavioral functions (Jennifer et al., 2012). The central clock orchestrates a common rhythm across the whole organism by synchronising with peripheral clocks including cardiovascular, metabolic, endocrine, immune and reproductive systems (Richards and Gumz, 2012). Lastly, these circadian oscillations can be entrained or modified by external inputs such as feeding behavior, social cues, temperature, and light (Rosenwasser and Turek, 2015).

### 1.2 Circadian terminology

Modelling circadian biology requires the use of sinosoid curves, characterised by mathematical properties unique to oscillating curves (illustrated in figure 1). For biological measurements to be considered circadian, their rhythms has to display a period of 24 hours. A period is an interval of time between successive cyclic pattern. Each period can be characterised by its highest point, called the zenith or the peak, and its lowest point called the nadir or the trough. The Mesor corresponds to the mean value between the zenith and the nadir of the oscillating curve. Mathematically, each period can be characterised by its amplitude and acrophase. The acrophase corresponds to the time where the curve reached its zenith for a given period, while the amplitude corresponds to the distance from the mesor to the zenith.

**Figure 1:**
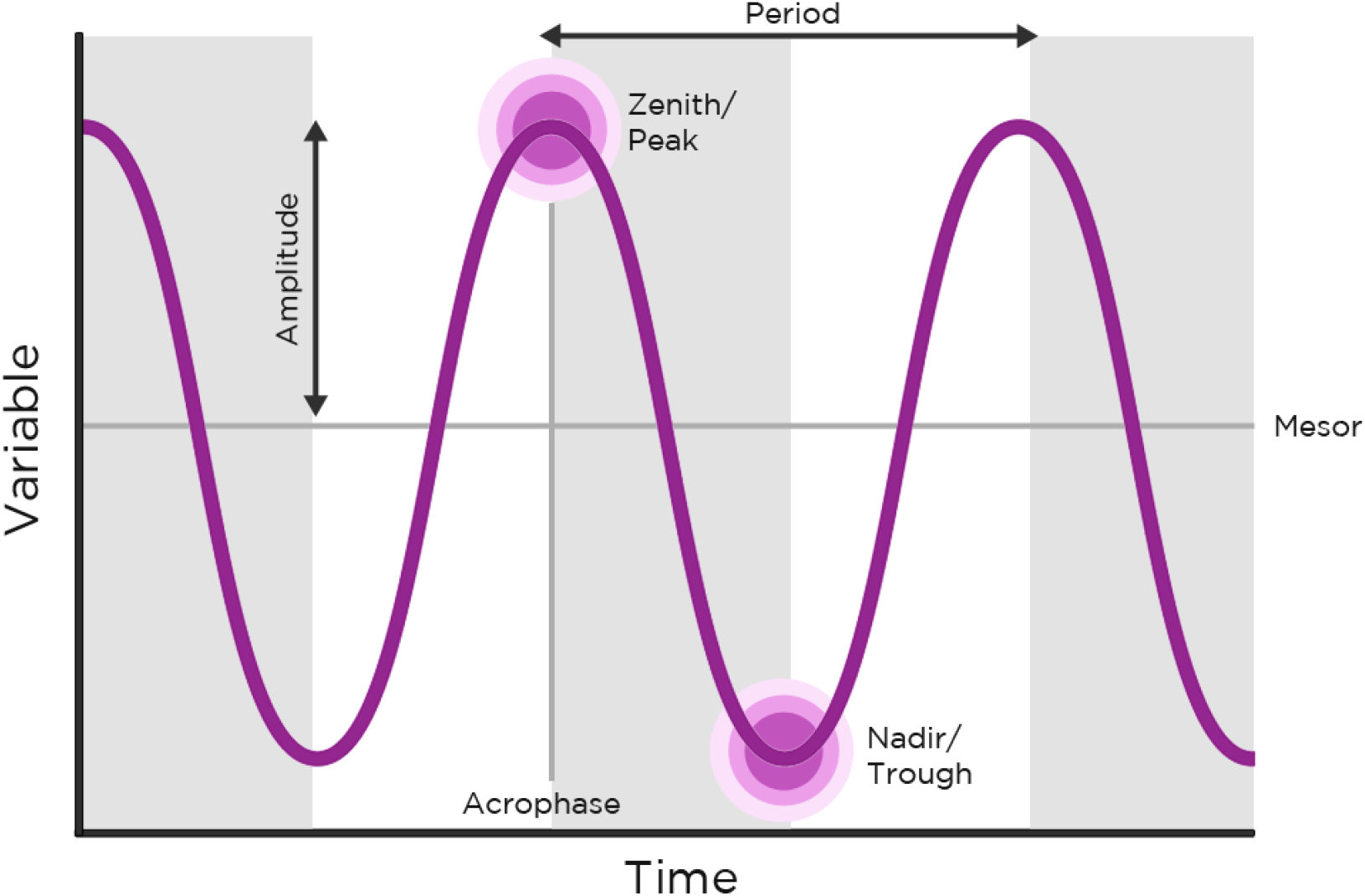
Graphical representation of a sinusoid curve indicating its properties of interest in circadian biology.

### 1.3 Related work

Some of the available options to analyse circadian biological data include JTK Cycle, Cosinor, and Lymorhyde. JTK Cycle is an algorithm that was designed to identify rhythmicity in genome scale data sets focusing on increasing power in order to be able to handle low-amplitude variables and is able to assess period, phase and amplitude (Hughes et al., 2010). Cosinor focuses on the analysis of time-series data and is able to determine rhythmicity and parameters of variables of known periods (Cornelissen, 2014). Lastly, Lymorhyde is based in linear models and can be used to detect differential rhythmicity in transcriptome data (Singer and Hughey, 2019).

### 1.4 Kronos

We aimed to provide a flexible framework to assess rhythmicity in biological data that remained convenient and accessible to researchers with a modest proficiency in coding. Kronos was designed based on already available open-access tools mentioned previously, with the aim to develop a package that can handle multivariate data sets, using base R linear modeling syntax. Kronos aims at filling this gap while being easy-to-use and implement. Some of Kronos’ features include the analysis of two or more groups, and more complex models such as two-factor designs. On top of that, Kronos also is designed to be flexible, assuming independence of data across timepoints, but can be adjusted to repeated-measures. Lastly, it can analyze any number of variables, and easily generate figures, being fully compatible with ggplot2 architecture.

## 2 Methods

Kronos is fully implemented in the R programming language and, while fully compatible with tidyverse code, only relies on the popular ggplot2 plotting package and on R packages that are considered base, such as the methods and stats packages.

Kronos supports standard R formula notation and relies on a custom kronosOut S4 object structure with specific ‘getter’ functions to provide organised and publication ready output tables.

Similar to the approaches of Cosinor and LimoRhyde, Kronos decomposes the time variable to a sine and cosine component for a set period. For example, if the period were to be 24 hours, the two components would be *sinθ* and *cosθ* where 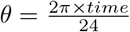.

Then, a simple generalised linear model can be fitted including both components like this:

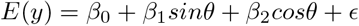

In this most basic case with a single set of observations without any factors or phenotypes to consider, rhythmicity can be assessed by assessing the explanatory potential of either the sine and/or cosine component of the model.

Similarly, in the case differential rhythmicity can be thought of as an interaction between the time components and an experimental variable or phenotype *P*, like this:

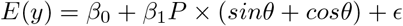

Which can be further expanded into:

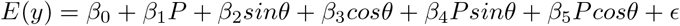

Here, differential rhythmicity can be thought of as the degree to which the terms *β*_4_*Psinθ* and *β*_5_*Pcosθ* explain the observed variance in the data in a manner not unlike an ANOVA.

## 3 Results

Kronos is implemented as a package in the R programming language (ver. 3.5+) and has been designed to equip circadian rhythm biologists with functions that can incorporate complex statistical designs while remaining user-friendly and streamlined, producing easily-interpreted data and helpful figures. For a complete description of the Kronos package functions please see our tutorial on Github: https://github.com/thomazbastiaanssen/kronos.

### 3.1 Input

Data should be prepared in “long form” for use in Kronos; that is with values repeating in the “Timepoint” column, which defines when data was collected during the period of interest (Wickham, 2014). Kronos has been designed and evaluated using data assuming a 24-hr period but has the flexibility to be applied to any period of interest. However, the period must be defined: Kronos does not have a function presently that can determine period length.

The package assumes data has been assessed for outliers and appropriately transformed.

We have included data sets with the package to demonstrate its features: it should be noted that the analysis performed here may not perfectly correspond to that performed in our published manuscripts. Furthermore, some variables in the data sets provided have been manipulated to better demonstrate functions of the package.

### 3.2 Estimating Rhythm Characteristics

Sinusoid curves can be predicted from each outcome variable using a user-defined period, which should be chosen by *a priori* hypothesis, or using other packages that allow for period detection. The kronos function produces a kronosOut object, providing the user with the following:

- The proportion of variance in the data explained by each individual predicted curve with the corresponding p-value along with the acrophase and amplitude.
- All the data required for graphing the sinusoid curve, which can either be used in our custom ggplot2 functions, or can be used in other graphing packages.
- Key details for the generated model that may be useful for prediction, modelling and other statistical applications.

### 3.3 Differential Rhythmicity

Kronos is designed to incorporate complex statistical designs and can be used to assess differential rhythmicity between two or more groups, and across o’mics data sets. The kronos function does this by running a generalised linear model on the categorical predictor variables given by the user, decomposed sine and cosine components of the data and their interactions. In these instances the kronosOut object contains additional information:

- The interaction between each independent factor included in the design and the overall time component. This is a composite score: the function collects all the p-values related to the factor of interest (as there are two time components), performs a Bonferroni’s adjustment and then returns the lowest resulting adjusted p-value.
- Pairwise comparisons for the model between all groups in order to assess differential rhythmicity.

This allows for the user to differentiate between differences overall and differences in rhythmicity between groups of interest. For example, in Figure 2A-B below both groups A and B exhibit rhythmicity (R^2^=0.219, p=0.031 and R^2^=0.331, p=0.003 respectively) and these rhythms are significantly different (F_1,56_=10.573, p=0.002). While group C does not exhibit rhythmicity (R^2^=0.023, p=0.723), overall the outcome variables is significantly elevated in group C relative to group A (F_1,56_=14.383, p<0.001).

**Figure 2:**
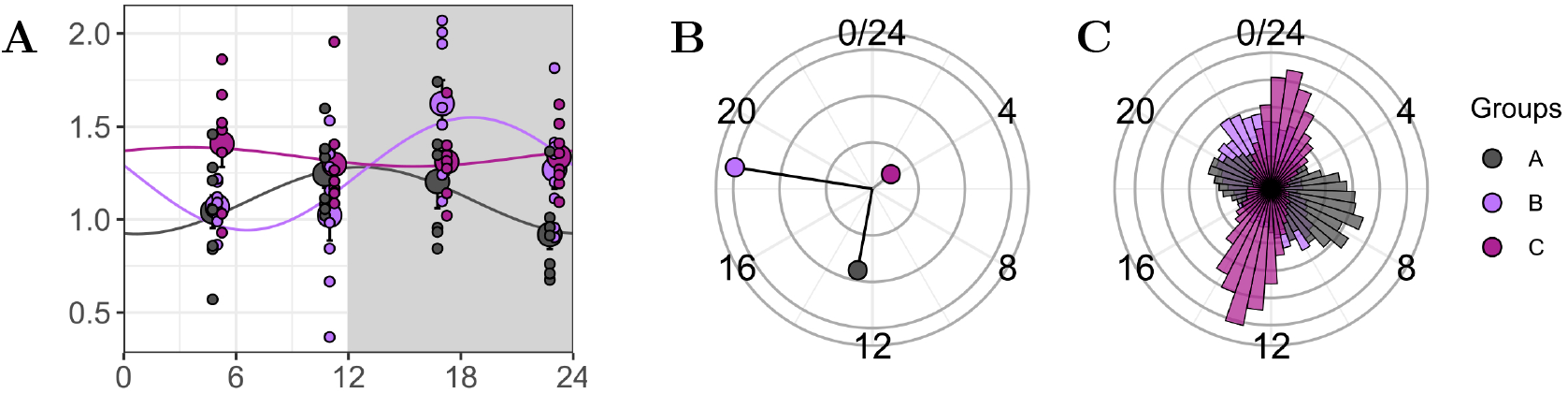
Differential rhythmic analysis output on mock data with three arbitrary experimental groups: A, B and C. A) A sinusoid curve representing the least-squares best fit trace per experimental group. Larger points represent groups means per time point, smaller points represent individual measurements. B) The acrophase and amplitude of the three groups assessed, represented on a circle plot for easy comparisons. Lengths of the lines represent amplitude normalised to the highest amplitude of that figure. Location on the circle depicts when during the cycle the acrophase takes place. Significantly rhythmic groups have an opaque line, whereas the line for group C, which does not reach significance for rhythmicity, is translucent. C) When working with high dimensional data, the kronos package can generate an acrogram, which shows the acrophase count data for all the variables in the data set. This allows researchers to easily identify temporal windows of biological significance in their data.

### 3.4 High dimensional/’omics data sets

Kronos can also be used to assess rhythmicity (both for individual groups and for more complex designs) in high dimensional or ‘omics data sets: the methods for this are laid out in detail in the Github tutorial. This approach can be used for transcriptomics, metabolomics, (meta-) proteomics and metagenomics - we initially built this package to assess circadian rhythms in the context of microbiota-gut-brain axis signalling. This is particularly useful as biological research has become increasingly reliant on these and other ‘omics approaches in recent years.

The kronosListToTable function generates easy-to-navigate statistical output for individual group rhythmicity, between-group rhythmicity and overall treatment effects. This function corrects for multiple comparisons automatically using base R FDR correction.

Additionally, gg_kronos_acrogram() can be utilised with omics data to represent group differences in spread of acrophases and amplitudes across a large data set (see Figure 2C).

### 3.5 Visualisation

Additionally the package contains custom ggplot2 figure functions, that utilise the kronos output to rapidly produce figures that convey important information for circadian rhythms (see Figure 2):

- gg_kronos_sinusoid() generates a x-y plot showing the outcome variable across the defined period. These graphs are useful for visualising the differences between specific timepoints assessed (Figure 2A).
- gg_kronos_circle() generates a plot showing the acrophase and amplitude of the predicted curve, allowing the reader to rapidly access summary data regarding variables of interest, and to compare the summary data between groups in more complex models. At baseline, non-significant outcome measures are presented using dashed lines (Figure 2B).
- gg_kronos_acrogram() generates a plot showing the acrophase for each feature in a high-dimensional data set, allowing the reader to rapidly access summary data of an entire data set, and to compare the summary data between groups in more complex models (Figure 2C).

### 3.6 Benchmarking against other commonly-used packages

In order to ensure Kronos performs similarly to other commonly-used packages, we benchmarked its performance against Cosinor2, JTK Cycle and LimoRhyde (for full details please see Supplementary Material or https://github.com/thomazbastiaanssen/kronos/blob/main/docs/benchmarking.md). We used a high dimensional data set containing one experimental group over time and assessed how each package identified significant rhythmicity. We found that Kronos and LimoRhyde performed most similarly, exhibiting a peak of p-values <0.05 and a uniform distribution >0.05 to 1. This distribution of p-values aligns with the preferred distribution of p-values for high dimensional data. Contrastingly, JTK Cycle exhibited a similar peak of p-values <0.05, but also contained another p-value peak at 0.95. Cosinor2 classified almost double the features as p-value < 0.05 compared to all other packages and the remaining p-values did not exhibit a uniform distribution.

When examining overlap in terms of features identified as rhythmic, Kronos did not identify any features that were not considered rhythmic by the other packages. In contrast, JTK Cycle identified 3 unique features, LimoRhyde identified 1 and Cosinor2 identified 303 additional unique features (see Figure 3).

**Figure 3:**
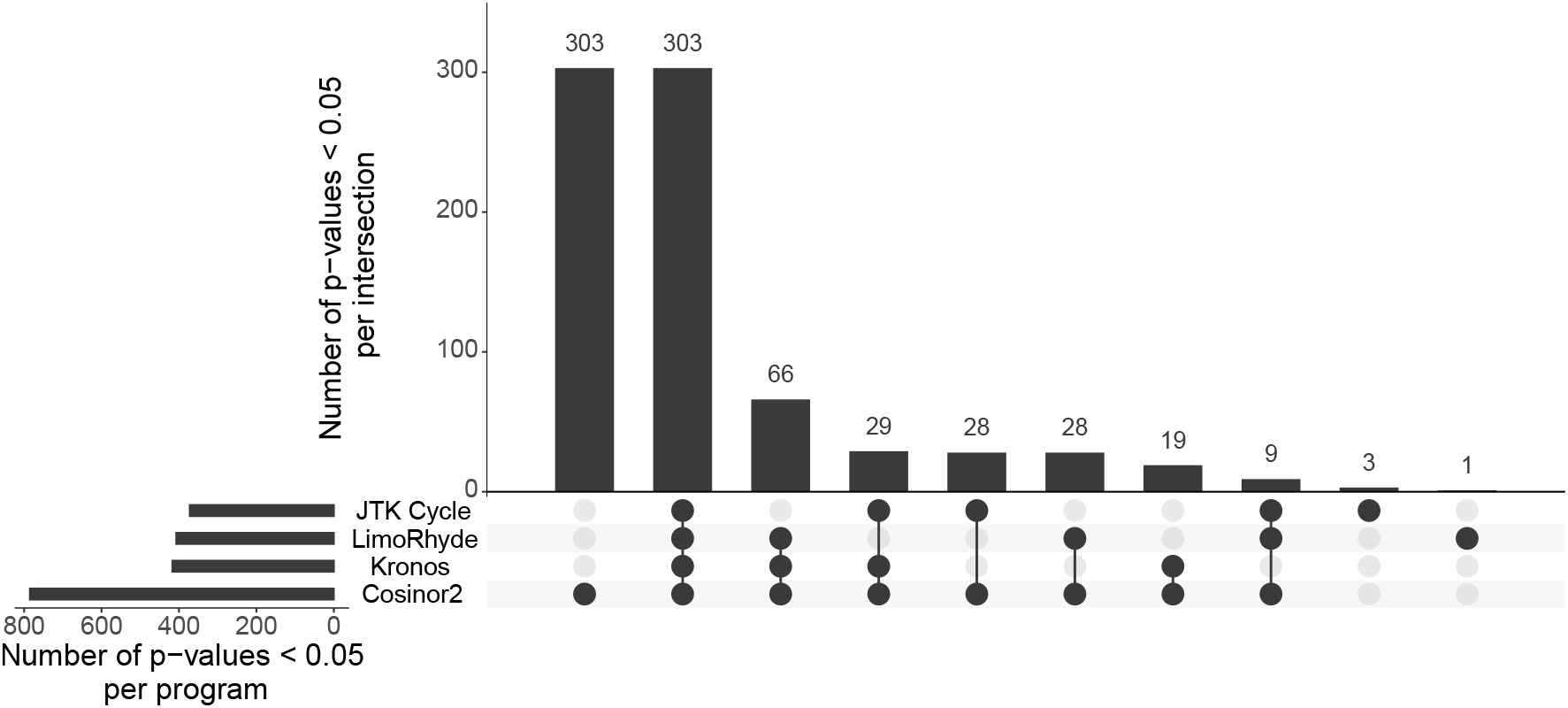
UpSet plot showing the number of features found to display rhythmicity at the p < 0.05 level as well as the degree of overlap between those features found by kronos and three commonly used programs: JTK Cycle, LimoRhyde and Cosinor2.

We then assessed whether the acrophases identified displayed concordance between packages. Here we did not compare LimoRhyde with with other packages as to our knowledge LimoRhyde does not provide raw acrophase values. We found that Kronos, JTK Cycle and Cosinor2 all exhibited concordance in acrophase detection (see Supplementary Material).

In summary, Kronos is a suitable method for assessing rhythmicity that does not sacrifice accuracy or reliability. It performs similarly to JTK Cycle and LimoRhyde in relation to significant p-value identification and overall p-value distribution in a high dimensional data set. Additionally, it produces concordant acrophases when compared to JTK Cycle and Cosinor2, indicating that it is performing to a similar standard.

## 4 Discussion

Kronos was designed to provide a flexible framework to assess circadian rhythms in biological data while being convenient and accessible to researchers with a modest proficiency in coding. Several other excellent packages exist to assess rhythmicity, but we found these to be more demanding both in terms of coding proficiency or unable to assess differential rhythmicity between groups. Kronos supports the standard R linear modeling formula syntax and comes with several publication-grade figure options out of the box. It also comes with a detailed and heavily annotated tutorial, preparing the researcher for independent analysis.

Nevertheless, Kronos still has numerous limitations. Most notably, Kronos has no means to estimate the period of a rhythm - the user is expected to have a predefined period as part of their hypothesis. Further, Kronos is unable to assess differential rhythmicity based on a continuous variable, only categorical variables are supported for this. Additionally, since Kronos decomposes time into exactly two components, it assumes that circadian rhythms are sinusoid and thus symmetrical.

Understanding chronobiology is essential for many different fields, but the data analysis involved is demanding. With the development and release of Kronos we set out to simplify the data analysis required. It is our hope that this will enable researchers from a variety of fields to study chronobiology in a more unencumbered and effective manner.

## Supporting information

Supplementary_benchmarking_analysis

## 5 Author Contributions

TFSB and SJL designed the package and wrote the tutorial with consultation from GSST and CEG. TFSB, SJL, GSST, CEG wrote the manuscript with input from all authors. SJL, GSST and CEG generated the biological data used to demonstrate the package. All authors provided critical feedback and helped shape the package, analysis and manuscript.

## 6 Acknowledgements

APC Microbiome Ireland is a research centre funded by Science Foundation Ireland (SFI), through the Irish Governments’ national development plan (grant no. 12/RC/2273_P2). Experiments involving circadian rhythms were in part funded by the Saks Kavanaugh Foundation. CEG is supported by European Foundation for the Advancement in Neurosciences, Geneva, Switzerland. SJL is funded by an Irish Research Council Postdoctoral Fellowship (GOPID/2021/298). The funding sources did not influence or constrain the study design, the collection, analysis, and interpretation of data, or the writing of the manuscript. We are grateful for the helpful discussions with Jorge Ricardo Soliz Rueda.

## 7 Declarations of Interest

JFC has been an invited speaker at conferences organized by Mead Johnson, Ordesa, and Yakult, and has received research funding from Reckitt, Nutricia, Dupont/IFF, and Nestle. GC has received honoraria from Janssen, Probi, and Apsen as an invited speaker; is in receipt of research funding from Pharmavite and Fonterra; and is a paid consultant for Yakult and Zentiva. This support neither influenced nor constrained the contents of this preview. TFSB, SJL, GSST, CEG, declare no competing interests.

## 8 Data and Code Availability

All data and code presented in this manuscript is open source and freely available at https://github.com/thomazbastiaanssen/kronos/ under the GPL-3 licence. Statistics are computed using the base R stats package.

## Notes

https://github.com/thomazbastiaanssen/kronos

