## Supplementary_benchmarking_analysis for "Kronos: A computational tool to facilitate biological rhythmicity analysis"

### Supplementary Materials: Benchmarking Kronos and other circadian packages

Thomaz F.S. Bastiaanssen

#### Preparation

Here, we will compare the performance of Kronos to that of three common R-based tools to assess rhythmic data. All code available at https://github.com/thomazbastiaanssen/kronos/blob/main/benchmark/

First, we will load the required packages.

```
#Wrangling
library(tidyverse)

#Plotting
library(ggforce)
library(ggvenn)
library(UpSetR)

#Load circadian rhythm packages
library(kronos) #devtools::install_github("thomazbastiaanssen/kronos")
library(limorhyde)
library(cosinor)
library(cosinor2)
```

Now we will load and prepare our high dimensional microbiome data for assessment.

```
source("00_load_and_clean_WGS.R")
```

Next, we apply the four packages to the data. Code to run JTK_CYCLE, Limorhyde and Cosinor2 was adapted from their respective vignettes.

```
source("01a_run_kronos.R")
source("01b_run_jtk_cycle.R")
source("01c_run_limorhyde.R")
source("01d_run_cosinor2.R")
```

Here, we collate all outcome parameters into a single object to compare them.

```
res_tot = do.call(cbind, list(res_kronos, res_jtk, res_limo, res_cosinor2))

colnames(res_tot) = paste(colnames(res_tot), c(rep("kronos", ncol(res_kronos)),
                                               rep("jtk",    ncol(res_jtk)),
                                               rep("limo",    ncol(res_limo)),
                                               rep("cosinor2",    ncol(res_cosinor2))),
                          sep = "_")
```

#### Comparing p-value distributions between programs

Theoretically, we expect a uniform distribution between 0-1 (for the cases where $H_0$ is true) as well as an inflation of low p-values (for the cases where $H_1$ is true).

```
par(mfcol = c(2, 2))

hist(res_kronos$p.val,   breaks = 20)
hist(res_jtk$p.val,      breaks = 20)
hist(res_cosinor2$p.val, breaks = 20)
hist(res_limo$p.val,     breaks = 20)
mtext("P-value histograms", side = 3, line = -1, outer = TRUE)
```

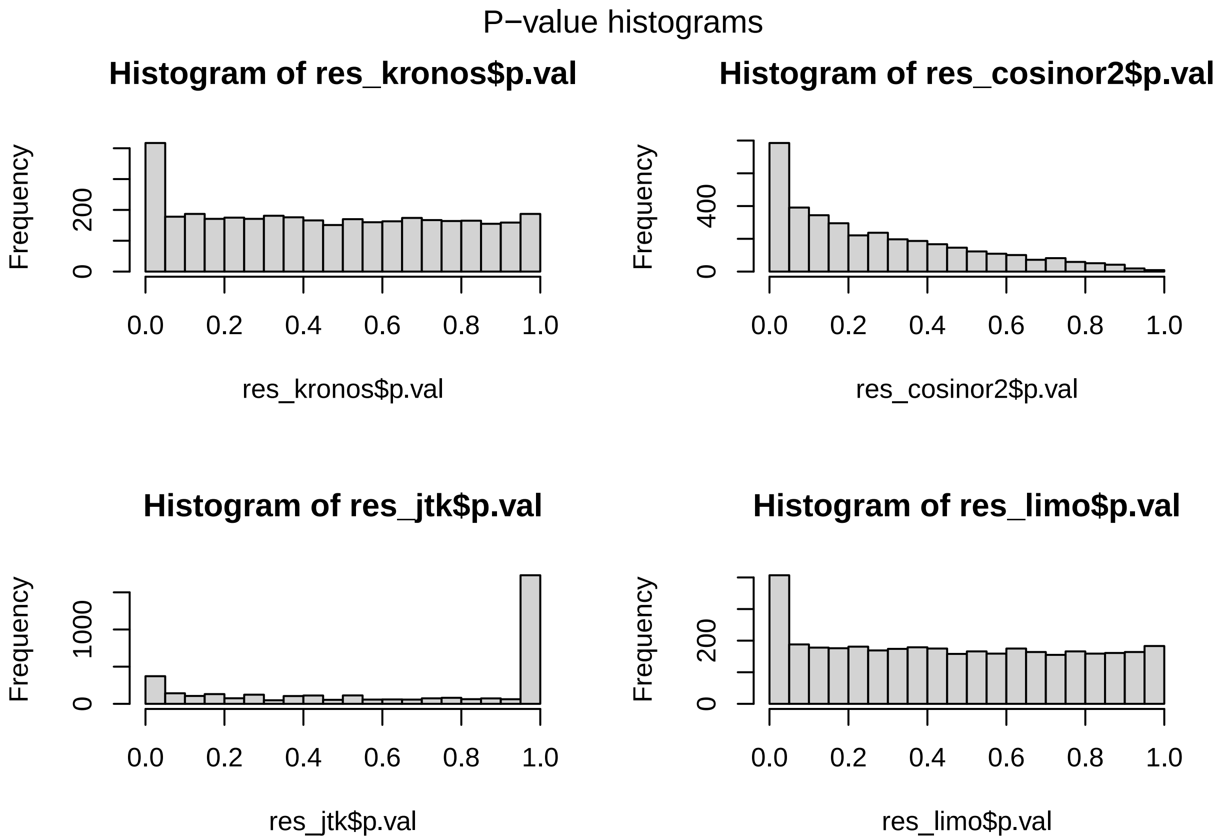


Both Limorhyde and Kronos show a desirable distribution of p-values, whereas JTK_CYCLE has an inflation on the higher end and Cosinor2 has a skewed, non-uniform distribution in the $H_0$ region.

#### Comparing acrophases between programs

```
res_tot %>%
  rownames_to_column("ID") %>%
  dplyr::select(!gene_id_limo) %>%
  group_by(ID) %>%
  filter(min(p.val_kronos,p.val_limo,p.val_jtk,p.val_cosinor2) < 0.05) %>%
  ungroup() %>%

  ggplot() +
  aes(x = .panel_x, y = .panel_y) +
```

```
  geom_point(aes(x = .panel_x, y = .panel_y)) +
  ggforce::facet_matrix(vars(acro_kronos, acro_jtk, acro_cosinor2)) +
  theme_bw() +
  ggtitle("Concordance in acrophase detection between Kronos, JTK_CYCLE and Cosinor2")
```

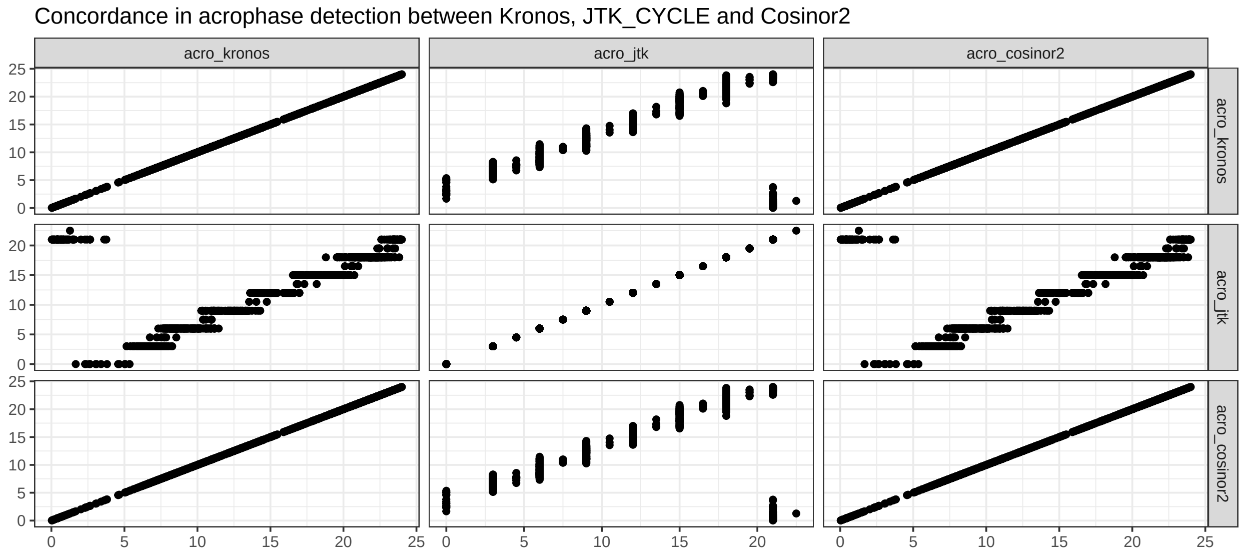


All three programs that produce acrophases are in agreement as to where the acrophases of features foudn to be rhythmic at $p < 0.05$ at least once are. Limorhyde does not have a clear method to return acrophases.

#### Plot agreement in hits between programs

In the venn diagram we see how many features are considered hits ($p < 0.05$) by the respective programs Alternatively, the UpSetR plot below shows the same.

```
venn_plot <- list(kronos    = row.names(res_tot[res_tot$p.val_kronos   < 0.05,]),
     jtk_cycle = row.names(res_tot[res_tot$p.val_jtk      < 0.05,]),
     limorhyde = row.names(res_tot[res_tot$p.val_limo     < 0.05,]),
     cosinor2  = row.names(res_tot[res_tot$p.val_cosinor2 < 0.05,])) %>%
  ggvenn::ggvenn(text_size = 2.5) +
  ggtitle("Venn diagram showing agreement between\nfeatures rhythmic at p < 0.05")

venn_plot$layers[[3]]$data$x <- c(-1, -0.8, 0.8, 1) #adjust margins
venn_plot
```

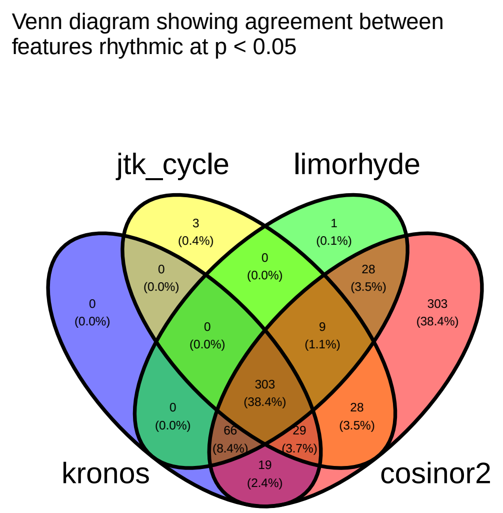


```r
list(kronos    = row.names(res_tot[res_tot$p.val_kronos   < 0.05,]),
     jtk_cycle = row.names(res_tot[res_tot$p.val_jtk      < 0.05,]),
     limorhyde = row.names(res_tot[res_tot$p.val_limo     < 0.05,]),
     cosinor2  = row.names(res_tot[res_tot$p.val_cosinor2 < 0.05,])) %>%
  UpSetR::fromList() %>%
  UpSetR::upset(., order.by = "freq",
                mainbar.y.label = "Number of p-values < 0.05\n per intersection",
                sets.x.label = "Number of p-values < 0.05\n per program")
```

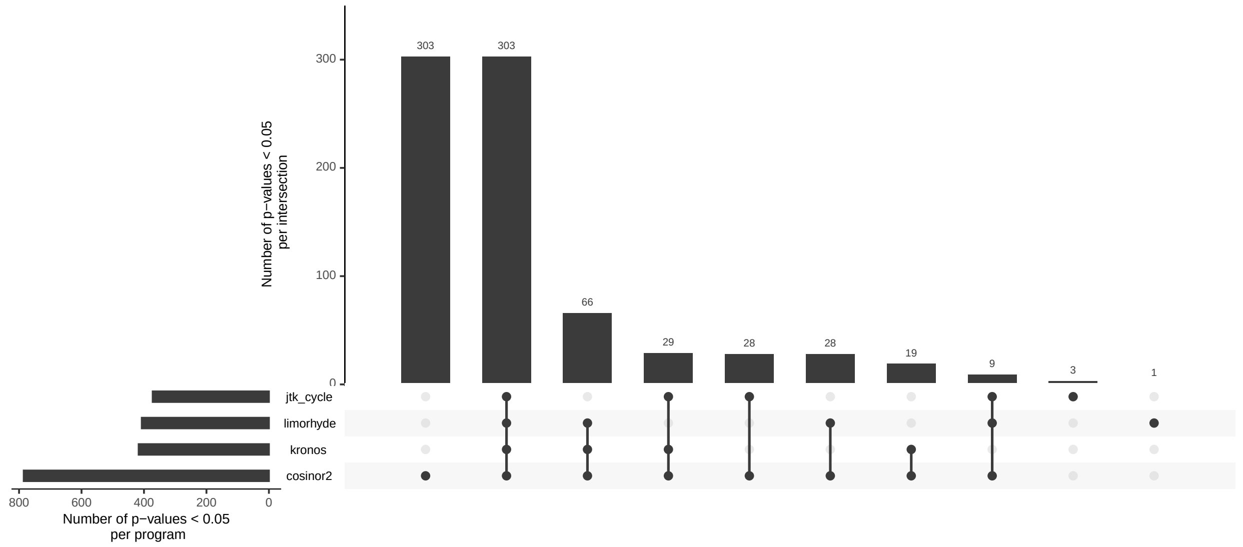


We can see that Kronos never reports a significntly rhythmic feature that at least one other program doesn't also see. Furthermore, Kronos finds the second highest total number of rhythmic features at p < 0.05. Only Cosinor2 finds more hits that Kronos, but 303 of those are not corroborated by any other tool and thus may be spurious.

From this, we can conclude that Kronos performs as well as (or better than) the best comparable program in each category. Furthermore, Kronos allows for convenient testing of arbitrarily complex models and comes with pre-designed ggplot-compatible plotting functions.

#### Session Info

```
sessioninfo::session_info()
```

```
## - Session info ---------------------------------------------------------------
##  setting  value
##  version  R version 4.2.2 Patched (2022-11-10 r83330)
##  os       Ubuntu 18.04.6 LTS
##  system   x86_64, linux-gnu
##  ui       X11
##  language en_IE:en
##  collate  en_IE.UTF-8
##  ctype    en_IE.UTF-8
##  tz       Europe/Dublin
##  date     2023-04-19
##  pandoc   2.19.2 @ /usr/lib/rstudio/resources/app/bin/quarto/bin/tools/ (via rmarkdown)
##
## - Packages -------------------------------------------------------------------
##  package          * version   date (UTC) lib source
##  annotate         * 1.74.0    2022-04-26 [1] Bioconductor
##  AnnotationDbi    * 1.58.0    2022-04-26 [1] Bioconductor
##  assertthat         0.2.1     2019-03-21 [1] CRAN (R 4.2.0)
##  backports          1.4.1     2021-12-13 [1] CRAN (R 4.2.0)
##  base64enc          0.1-3     2015-07-28 [1] CRAN (R 4.2.0)
##  Biobase          * 2.56.0    2022-04-26 [1] Bioconductor
##  BiocGenerics     * 0.42.0    2022-04-26 [1] Bioconductor
##  Biostrings         2.64.1    2022-08-18 [1] Bioconductor
##  bit                4.0.5     2022-11-15 [1] CRAN (R 4.2.1)
##  bit64              4.0.5     2020-08-30 [1] CRAN (R 4.2.0)
##  bitops             1.0-7     2021-04-24 [1] CRAN (R 4.2.0)
##  blob               1.2.3     2022-04-10 [1] CRAN (R 4.2.0)
##  broom              1.0.2     2022-12-15 [1] CRAN (R 4.2.1)
##  cachem             1.0.6     2021-08-19 [1] CRAN (R 4.2.0)
##  cellranger         1.1.0     2016-07-27 [1] CRAN (R 4.2.0)
##  checkmate          2.1.0     2022-04-21 [1] CRAN (R 4.2.0)
##  cli                3.6.0     2023-01-09 [1] CRAN (R 4.2.1)
##  cluster            2.1.4     2022-08-22 [4] CRAN (R 4.2.1)
##  codetools          0.2-19    2023-02-01 [4] CRAN (R 4.2.2)
##  colorspace         2.0-3     2022-02-21 [1] CRAN (R 4.2.0)
##  cosinor          * 1.2.2     2022-05-24 [1] CRAN (R 4.2.2)
##  cosinor2         * 0.2.1     2018-10-15 [1] CRAN (R 4.2.0)
##  cowplot            1.1.1     2020-12-30 [1] CRAN (R 4.2.0)
##  crayon             1.5.2     2022-09-29 [1] CRAN (R 4.2.1)
##  data.table       * 1.14.6    2022-11-16 [1] CRAN (R 4.2.1)
##  DBI                1.1.3     2022-06-18 [1] CRAN (R 4.2.0)
##  dbplyr             2.3.0     2023-01-16 [1] CRAN (R 4.2.1)
##  deldir             1.0-6     2021-10-23 [1] CRAN (R 4.2.1)
##  deleuze          * 0.1.0.0   2023-01-11 [1] Github (thomazbastiaanssen/deleuze@129f472)
##  digest             0.6.31    2022-12-11 [1] CRAN (R 4.2.1)
##  dplyr            * 1.0.10    2022-09-01 [1] CRAN (R 4.2.1)
##  ellipsis           0.3.2     2021-04-29 [1] CRAN (R 4.2.0)
##  evaluate           0.20      2023-01-17 [1] CRAN (R 4.2.1)
##  fansi              1.0.3     2022-03-24 [1] CRAN (R 4.2.0)
##  farver             2.1.1     2022-07-06 [1] CRAN (R 4.2.1)
```

```
##  fastmap            1.1.0    2021-01-25 [1] CRAN (R 4.2.0)
##  forcats          * 0.5.2    2022-08-19 [1] CRAN (R 4.2.1)
##  foreign            0.8-82   2022-01-13 [4] CRAN (R 4.1.2)
##  Formula            1.2-4    2020-10-16 [1] CRAN (R 4.2.0)
##  fs                 1.5.2    2021-12-08 [1] CRAN (R 4.2.0)
##  gargle             1.2.1    2022-09-08 [1] CRAN (R 4.2.1)
##  genefilter         1.78.0   2022-04-26 [1] Bioconductor
##  generics           0.1.3    2022-07-05 [1] CRAN (R 4.2.1)
##  GenomeInfoDb       1.32.4   2022-09-06 [1] Bioconductor
##  GenomeInfoDbData   1.2.8    2022-05-09 [1] Bioconductor
##  ggforce          * 0.4.1    2022-10-04 [1] CRAN (R 4.2.1)
##  ggplot2          * 3.4.0    2022-11-04 [1] CRAN (R 4.2.1)
##  ggvenn           * 0.1.9    2021-06-29 [1] CRAN (R 4.2.2)
##  glue               1.6.2    2022-02-24 [1] CRAN (R 4.2.0)
##  googledrive        2.0.0    2021-07-08 [1] CRAN (R 4.2.0)
##  googlesheets4      1.0.1    2022-08-13 [1] CRAN (R 4.2.1)
##  gridExtra          2.3      2017-09-09 [1] CRAN (R 4.2.0)
##  gtable             0.3.1    2022-09-01 [1] CRAN (R 4.2.1)
##  haven              2.5.1    2022-08-22 [1] CRAN (R 4.2.1)
##  highr              0.10     2022-12-22 [1] CRAN (R 4.2.1)
##  Hmisc              4.7-2    2022-11-18 [1] CRAN (R 4.2.1)
##  hms                1.1.2    2022-08-19 [1] CRAN (R 4.2.1)
##  htmlTable          2.4.1    2022-07-07 [1] CRAN (R 4.2.1)
##  htmltools          0.5.4    2022-12-07 [1] CRAN (R 4.2.1)
##  htmlwidgets        1.6.1    2023-01-07 [1] CRAN (R 4.2.1)
##  httpuv             1.6.8    2023-01-12 [1] CRAN (R 4.2.1)
##  httr               1.4.4    2022-08-17 [1] CRAN (R 4.2.1)
##  interp             1.1-3    2022-07-13 [1] CRAN (R 4.2.1)
##  IRanges          * 2.30.1   2022-08-18 [1] Bioconductor
##  jpeg               0.1-10   2022-11-29 [1] CRAN (R 4.2.1)
##  jsonlite           1.8.4    2022-12-06 [1] CRAN (R 4.2.1)
##  KEGGREST           1.36.3   2022-07-12 [1] Bioconductor
##  knitr              1.41     2022-11-18 [1] CRAN (R 4.2.1)
##  kronos           * 0.1.0.0  2023-03-08 [1] Github (thomazbastiaanssen/kronos@4e9a18c)
##  labeling           0.4.2    2020-10-20 [1] CRAN (R 4.2.0)
##  later              1.3.0    2021-08-18 [1] CRAN (R 4.2.0)
##  lattice            0.20-45  2021-09-22 [4] CRAN (R 4.2.0)
##  latticeExtra       0.6-30   2022-07-04 [1] CRAN (R 4.2.1)
##  lifecycle          1.0.3    2022-10-07 [1] CRAN (R 4.2.1)
##  limma            * 3.52.4   2022-09-27 [1] Bioconductor
##  limorhyde        * 1.0.1    2022-02-18 [1] CRAN (R 4.2.2)
##  lubridate          1.9.0    2022-11-06 [1] CRAN (R 4.2.1)
##  magrittr           2.0.3    2022-03-30 [1] CRAN (R 4.2.0)
##  MASS               7.3-58.2 2023-01-23 [4] CRAN (R 4.2.2)
##  Matrix             1.5-3    2022-11-11 [1] CRAN (R 4.2.1)
##  matrixStats        0.63.0   2022-11-18 [1] CRAN (R 4.2.1)
##  memoise            2.0.1    2021-11-26 [1] CRAN (R 4.2.0)
##  mime               0.12     2021-09-28 [1] CRAN (R 4.2.0)
##  modelr             0.1.10   2022-11-11 [1] CRAN (R 4.2.1)
##  munsell            0.5.0    2018-06-12 [1] CRAN (R 4.2.0)
##  nnet               7.3-18   2022-09-28 [4] CRAN (R 4.2.1)
##  pillar             1.8.1    2022-08-19 [1] CRAN (R 4.2.1)
##  pkgconfig          2.0.3    2019-09-22 [1] CRAN (R 4.2.0)
##  plyr               1.8.8    2022-11-11 [1] CRAN (R 4.2.1)
```

```
##  png               0.1-8     2022-11-29 [1] CRAN (R 4.2.1)
##  polyclip          1.10-4    2022-10-20 [1] CRAN (R 4.2.1)
##  promises          1.2.0.1   2021-02-11 [1] CRAN (R 4.2.0)
##  purrr           * 1.0.1     2023-01-10 [1] CRAN (R 4.2.1)
##  R6                2.5.1     2021-08-19 [1] CRAN (R 4.2.0)
##  RColorBrewer      1.1-3     2022-04-03 [1] CRAN (R 4.2.0)
##  Rcpp              1.0.9     2022-07-08 [1] CRAN (R 4.2.1)
##  RCurl             1.98-1.9  2022-10-03 [1] CRAN (R 4.2.1)
##  readr           * 2.1.3     2022-10-01 [1] CRAN (R 4.2.1)
##  readxl            1.4.1     2022-08-17 [1] CRAN (R 4.2.1)
##  reprex            2.0.2     2022-08-17 [1] CRAN (R 4.2.1)
##  rlang             1.0.6     2022-09-24 [1] CRAN (R 4.2.1)
##  rmarkdown         2.20      2023-01-19 [1] CRAN (R 4.2.1)
##  rpart             4.1.19    2022-10-21 [4] CRAN (R 4.2.1)
##  RSQLite           2.2.20    2022-12-22 [1] CRAN (R 4.2.1)
##  rstudioapi        0.14      2022-08-22 [1] CRAN (R 4.2.1)
##  rvest             1.0.3     2022-08-19 [1] CRAN (R 4.2.1)
##  S4Vectors       * 0.34.0    2022-04-26 [1] Bioconductor
##  scales            1.2.1     2022-08-20 [1] CRAN (R 4.2.1)
##  sessioninfo       1.2.2     2021-12-06 [1] CRAN (R 4.2.0)
##  shiny             1.7.4     2022-12-15 [1] CRAN (R 4.2.1)
##  stringi           1.7.12    2023-01-11 [1] CRAN (R 4.2.1)
##  stringr         * 1.5.0     2022-12-02 [1] CRAN (R 4.2.1)
##  survival          3.4-0     2022-08-09 [4] CRAN (R 4.2.1)
##  tibble          * 3.1.8     2022-07-22 [1] CRAN (R 4.2.1)
##  tidyr           * 1.2.1     2022-09-08 [1] CRAN (R 4.2.1)
##  tidyselect        1.2.0     2022-10-10 [1] CRAN (R 4.2.1)
##  tidyverse       * 1.3.2     2022-07-18 [1] CRAN (R 4.2.1)
##  timechange        0.2.0     2023-01-11 [1] CRAN (R 4.2.1)
##  tweenr            2.0.2     2022-09-06 [1] CRAN (R 4.2.1)
##  tzdb              0.3.0     2022-03-28 [1] CRAN (R 4.2.0)
##  UpSetR          * 1.4.0     2019-05-22 [1] CRAN (R 4.2.2)
##  utf8              1.2.2     2021-07-24 [1] CRAN (R 4.2.0)
##  vctrs             0.5.1     2022-11-16 [1] CRAN (R 4.2.1)
##  withr             2.5.0     2022-03-03 [1] CRAN (R 4.2.0)
##  xfun              0.36      2022-12-21 [1] CRAN (R 4.2.1)
##  XML             * 3.99-0.13 2022-12-04 [1] CRAN (R 4.2.1)
##  xml2              1.3.3     2021-11-30 [1] CRAN (R 4.2.0)
##  xtable            1.8-4     2019-04-21 [1] CRAN (R 4.2.0)
##  XVector           0.36.0    2022-04-26 [1] Bioconductor
##  yaml              2.3.6     2022-10-18 [1] CRAN (R 4.2.1)
##  zlibbioc          1.42.0    2022-04-26 [1] Bioconductor
##
##  [1] /home/thomaz/R/x86_64-pc-linux-gnu-library/4.2
##  [2] /usr/local/lib/R/site-library
##  [3] /usr/lib/R/site-library
##  [4] /usr/lib/R/library
##
## ------------------------------------------------------------------------------
```
